# Impact of Organic Compounds on the Stability of Influenza A Virus in deposited 1-µl droplets

**DOI:** 10.1101/2024.05.20.595000

**Authors:** Aline Schaub, Shannon C. David, Irina Glas, Liviana K. Klein, Kalliopi Violaki, Céline Terrettaz, Ghislain Motos, Nir Bluvshtein, Beiping Luo, Marie Pohl, Walter Hugentobler, Athanasios Nenes, Silke Stertz, Ulrich K. Krieger, Thomas Peter, Silke Stertz, Tamar Kohn

## Abstract

The composition of respiratory fluids influences the stability of viruses in exhaled aerosol particles and droplets, though the role of respiratory organics in modulating virus stability remains poorly understood. This study investigates the effect of organic compounds on the stability of influenza A virus (IAV) in deposited droplets. We compare the infectivity loss of IAV at different relative humidities (RH) over the course of one hour in 1-μl droplets consisting of phosphate-buffered saline (without organics), synthetic lung fluid, or nasal mucus (both containing organics). We show that IAV stability increases with increasing organic:salt ratios. Among the various organic species, proteins are identified as the most protective component, with smaller proteins stabilizing IAV more efficiently at the same mass concentration. Organics act by both increasing the efflorescence RH and shortening the drying period until efflorescence at a given RH. This research advances our mechanistic understanding of how organics stabilize exhaled viruses and thus influence their inactivation in respiratory droplets.

## Introduction

Influenza A virus (IAV) is a respiratory pathogen transmitted in part via the airborne route through exhaled aerosol particles and droplets.^1–5^ Airborne transmission necessitates that IAV remains infectious within particles exhaled by an infected individual until they are inhaled by a new host. These aerosol particles and droplets are composed of a complex mixture of salts and organic compounds.^6– 11^ The exact composition depends on the origin of the particle within the host^6,12,13^ (nose, mouth, or upper or lower respiratory tract), and compositions have been shown to differ between individuals.^12^ Moreover, the physical condition of an individual can also modulate the composition of the fluid, with a higher mucus content observed when a person is infected by IAV.^14^

After exhalation, the aerosol particles and droplets encounter typical indoor air conditions (room temperature, RH ≈ 30-50%,^15^ ambient trace gases), which differ dramatically from those encountered in the respiratory tract with respect to temperature (≈ 37°C), RH (≈ 95%) and gas composition. This leads to changes in the microenvironment surrounding the virus, namely rapid evaporation of water, which is determined by the RH of the surrounding air. Consequently, solute concentrations increase and phase transitions may occur, including salt efflorescence,^16^ liquid-liquid phase separation,^17^ or the formation of an organic glassy shell.^8^ In addition, concentrations of CO_2_ and NH_4_ in the particles (which are known to be high upon exhalation) are reduced via equilibration with the indoor air, while trace gases from the ambient air are taken up by the particles. This internal micro-environment is therefore highly dynamic, and contributes to explain the observed dependence of IAV infectivity on the RH of indoor air.^18,19,28–30,20–27^

Among the physical-chemical parameters modulated by RH, pH^16^ and NaCl molality^31^ have been identified as important drivers for IAV inactivation in prior work. Past studies have also shown that organic biomolecules, such as those contained in respiratory fluids, stabilize the virus against these stressors.^18,19,32–35^ Specifically, Yang et al.^19^ and Kormuth et al.^18^ observed a dependence of IAV infectivity on RH in aerosol particles and deposited droplets, whereby the lowest infectivity was observed at intermediate RH. This RH dependence was suppressed when organic material, such as fetal calf serum or extracellular material from epithelial cell cultures, were added to the matrix. The specific organic components and the mechanisms responsible for this protection are unknown, but proteins are suspected as a possible cause. The protective role of proteins has been supported additional studies proposing which mechanisms to explain this effect: Huang et al.^34^ observed that an increase in protein concentration in a drying droplet modified its dried morphology by triggering the “coffee-ring effect”, i.e., the deposition of the droplet components at the droplet periphery. This resulted in the aggregation of viruses (bacteriophage Phi6, in their study) and proteins in the reduced volume of the ring, thereby reducing their exposure to physical-chemical stressors. A different mechanism was suggested by Huynh et al.^35^, who reported an increase in viscosity with increasing protein content, and proposed that the resulting semisolid phase in aerosol particles and deposited droplets protects the virus by limiting solute diffusion. Furthermore, non-proteinaceous organics were also identified to stabilize viruses. For example, Nieto-Caballero et al.^33^ observed the vitrification of carbohydrates present in saliva, and the resulting increase of viscosity protected murine hepatitis virus in aerosol particles, potentially by stabilizing the virus surface proteins. This is consistent with our previous work suggesting that a carbohydrate (in this case sucrose) contributes to the protection of IAV in saline droplets, most likely by stabilizing IAV surface proteins and lipid membranes.^31^ Additionally, sucrose lowers the Na^+^ and Cl^−^ molalities in the drying droplets, thereby slowing the inactivation of IAV by the salt. This latter effect is particularly pronounced immediately prior to efflorescence, when salt molalities in the droplets are at their highest and reach massive supersaturations. Moreover, protection of IAV against salt-mediated inactivation was observed in saline droplets containing commensal respiratory bacteria.^36^ Here, the bacteria were found to flatten the droplet and hence shorten the drying time until efflorescence occurs, and thereby reduce the exposure of IAV to deleterious, supersaturated NaCl solutions.

These studies emphasize the importance of matrix composition for exhaled virus stability. However, respiratory fluids are complex aqueous mixtures of salts, proteins, lipids, sugars, and surfactants,^7,8,37– 43^ and the effect of each component on virus stability remains to be assessed. Here, we quantified the infectivity of IAV in 1-μl deposited droplets at RH levels ranging from 15 to 95% in three matrices, namely phosphate-buffered saline (PBS), synthetic lung fluid (SLF), and nasal mucus collected from epithelial cells. Simultaneously, we monitored salt efflorescence in these droplets. To disentangle the role of different organics, we then decomposed SLF to test the importance of individual components. In addition, we assessed the virus-stabilizing function of proteins as a function of protein size and content.

## Materials and methods

### Virus propagation and purification

Madin-Darby Canine Kidney (MDCK) cells (ThermoFisher) maintained in Dulbecco’s modified Eagle medium (DMEM, Gibco) supplemented with 10% Fetal Bovine Serum (FBS; Gibco), and 1% Penicillin-Streptomycin 10,000 U/ml (P/S; Gibco) were used to propagate influenza A virus (IAV) strain A/WSN/33 (H1N1 subtype). Washed confluent MDCK cells were was inoculated with IAV at a multiplicity of infection (MOI) of 0.001 for 72 h in OptiMEM (Gibco) supplemented with 1% P/S and 1 μg/ml TPCK trypsin (Sigma, T1426). Centrifugation (2,500 ×g, 10 min) was used to clear the culture supernatants from the infected cells. Ultracentrifugation (112,400 ×g in a SW31Ti rotor (Beckman), 90 min, 4°C) was subsequently used to concentrate the virus by pelleting through a 30% sucrose cushion. PBS (ThermoFisher, 18912014) was used to resuspend the pellets overnight at 4°C. After quantification by plaque assay (described below), the concentrated IAV stock solution was determined to be ∼10^10^ plaque forming units (PFU)/ml, and frozen at −80°C until use.

### Quantification of IAV infectivity

IAV infectivity was quantified by plaque assay on MDCK cells, as described in David et al.^44^ Briefly, cell monolayers were prepared in 12-well plates and were washed with PBS prior to infection with IAV samples. The samples were initially diluted in series using PBSi (PBS for infection; PBS supplemented with 1% P/S, 0.02 mM Mg^2+^, 0.01 mM Ca^2+^, and 0.3% bovine serum albumin (Sigma-Aldrich A1595); pH ∼7.3). PBSi was also used as a negative control on an extra plate. Infected cells were incubated for 1 h at 37°C with 5% of CO_2_, and manually agitated every 10 min. The remaining inoculum was then removed and an agar overlay (MEM supplemented with 0.5 μg/ml of TPCK-trypsin, 0.01% DEAE-dextran, 0.11% sodium bicarbonate, and 0.7% of Oxoid agar (Thermo Fisher LP0028-500G) was used to cover the cells. After 72 h of incubation (37°C, 5% CO_2_), the cells were fixed with 10% of formaldehyde (Sigma, 47608-1L-F) in PBS and stained with a 5% crystal violet solution (Sigma, HT901-8FOZ) in water and 10% methanol (Fisher Chemical, M-4000-15). Plaques were enumerated to determine the virus titer in PFU/ml. The limit of quantification (LOQ) of this assay was 10 PFU/ml.

### Quantification of IAV genome copies by RT-qPCR

The QIAamp Viral RNA Mini extraction kit (Qiagen, 52906) was used to perform RNA extractions, according to manufacturer’s instructions. 140 μl of each sample were extracted and 80 μl of elution buffer were used to recover extracted nucleic acids. The extracts were stored at −20°C until analysis. Water was extracted as a negative control and was always negative. A Mic Real-Time PCR System from Bio Molecular Systems was used to perform amplification and detection, with the One Step PrimeScript™ RT-PCR Kit (Takara Bio, RR064A). The following primers targeting a 244-base amplicon of the IAV M segment were used:

Forward primer 5’-ATGAGYCTTYTAACCGAGGTCGAAACG-3’

Reverse primer 5’-TGGACAAANCGTCTACGCTGCAG-3’

The reaction mixture used for the one-step RT-qPCR assay was composed of 7.5 μl of 2x One-Step SYBR RT-PCR Buffer, 0.3 μl of Takara Ex Taq HS (5 U/μl stock), 0.3 μl PrimeScript RT enzyme mix, 0.3 μl forward and reverse primers (10 μM stocks), and 3.3 μl RNase-free water, to which 3 μl extracted RNA were added. The run profile was the following: 2 min at 50°C and 10 min at 95°C for reverse transcription and denaturation, followed by 40 cycles of 15 sec at 95°C and 60 sec and 60°C for annealing and extension, and a final dissociation step from 55°C to 95°C at 0.3°C/s. A standard curve was created for quantification over a range of 10 to 10^8^ genomic copies (GC)/μl using a Gblock gene fragment, as described in ^45^. A no-template control of milli-Q water was included in every run and was always negative. Serial dilutions of the samples were used regularly to check for the absence of inhibition. The qPCR data were acquired with the micPCR software (version 2.12.16). Pooled standard curves were analyzed using the Generic qPCR Limit of Detection (LOD) / Limit of Quantification (LOQ) calculator ^46^. The average slope of the standard curve was −3.44 and the average intercept was 34.08, with an R^2^ of 0.99 and a PCR efficiency of 95.5%. The LOQ, defined here as the lowest standard concentration with a coefficient of variation smaller 35%, was determined at 10 copies/reaction. All RT-qPCR procedures followed MIQE guidelines^47^ (see **Table S1**).

### Matrix preparation

Experiments were conducted in three main matrices: PBS, SLF, and nasal mucus. PBS was bought as salt solution from ThermoFisher (18912014). SLF was prepared following the recipe adapted from Bicer^7^ as detailed in Luo et al.^16^ and was freeze-dried according to the method described by Hassoun et al.^42^ Constituents of SLF are Hank’s Balanced Salt Solution (HBSS) without phenol red, lyophilized albumin from human serum, human transferrin, 1,2-dipalmitoyl-sn-glycero-3-phosphocholine (DPPC), 1,2-dipalmitoyl-sn-glycero-3-phospho-rac-(1-glycerol) ammonium salt (DPPG), cholesterol, L-ascorbic acid, uric acid and glutathione, which were all purchased from Sigma-Aldrich. SLF powder was resuspended in milli-Q water. Decomposed SLF was prepared following the same method, but omitting the components of interest. Nasal mucus was harvested from primary epithelial nasal cultures (Epithelix, Switzerland, # EP51AB) grown at the air-liquid interface as described in Luo et al.^16^ Two batches were obtained, of which one was used for virus inactivation experiments and the other for composition analysis. The salt, protein, and lipid contents of mucus were measured as described in the Supporting Information. The composition of all three matrices is detailed in **Table 1**.

**Table 1:**
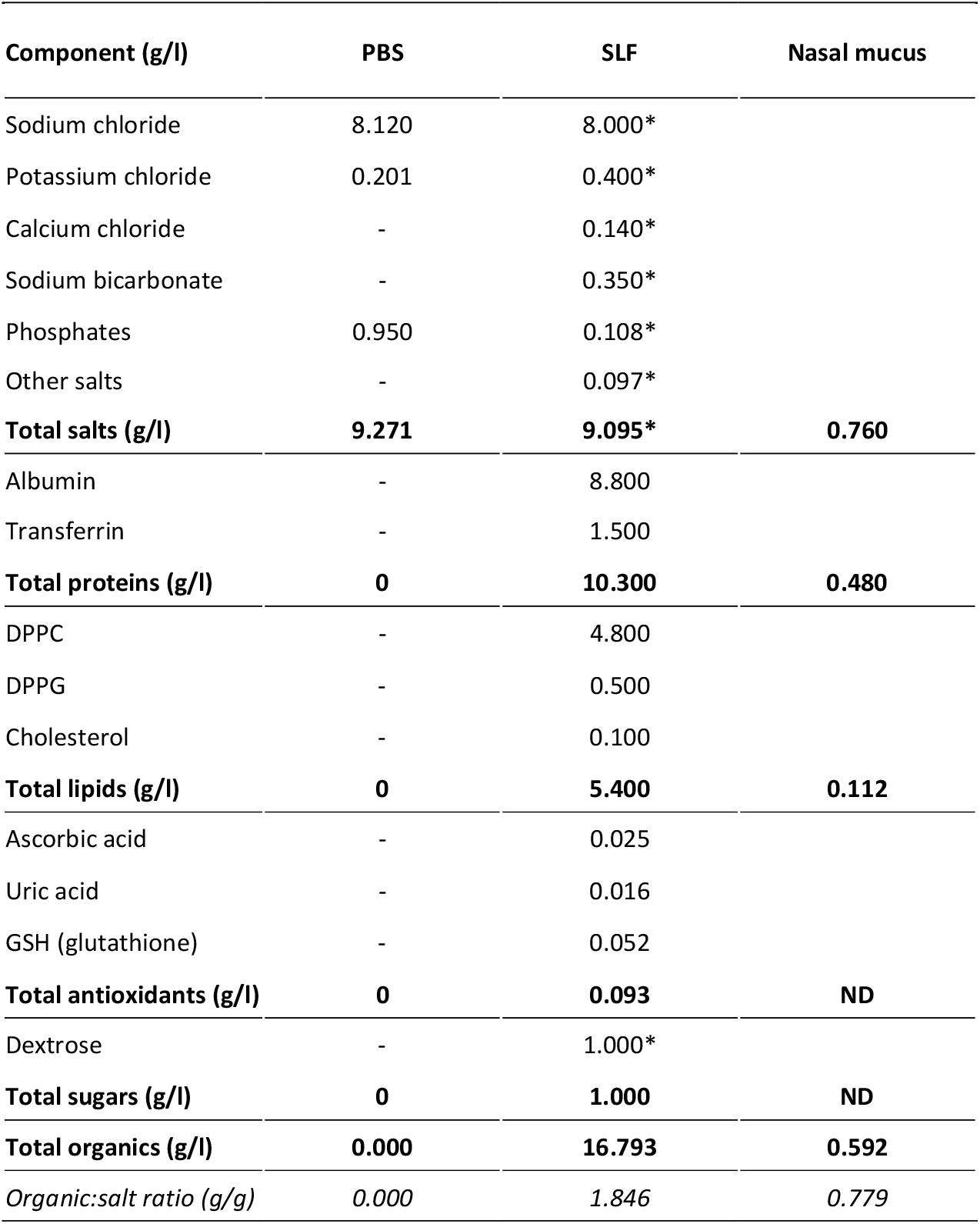
Composition of synthetic fluids and nasal mucus. Proteins, lipids, and antioxidants in SLF were dissolved in HBSS (components are indicated with an asterisk). In nasal mucus, only the total content of salts, proteins and lipids is listed. Further details on specific ions and lipids are provided in the SI (**Tables S2** and **S3**, respectively). ND = not determined.

| Component (g/l) | PBS | SLF | Nasal mucus |
| --- | --- | --- | --- |
| Sodium chloride | 8.120 | 8.000* |  |
| Potassium chloride | 0.201 | 0.400* |  |
| Calcium chloride | - | 0.140* |  |
| Sodium bicarbonate | - | 0.350* |  |
| Phosphates | 0.950 | 0.108* |  |
| Other salts | - | 0.097* |  |
| **Total salts (g/l)** | **9.271** | **9.095\*** | **0.760** |
| Albumin | - | 8.800 |  |
| Transferrin | - | 1.500 |  |
| **Total proteins (g/l)** | **0** | **10.300** | **0.480** |
| DPPC | - | 4.800 |  |
| DPPG | - | 0.500 |  |
| Cholesterol | - | 0.100 |  |
| **Total lipids (g/l)** | **0** | **5.400** | **0.112** |
| Ascorbic acid | - | 0.025 |  |
| Uric acid | - | 0.016 |  |
| GSH (glutathione) | - | 0.052 |  |
| **Total antioxidants (g/l)** | **0** | **0.093** | **ND** |
| Dextrose | - | 1.000* |  |
| **Total sugars (g/l)** | **0** | **1.000** | **ND** |
| **Total organics (g/l)** | **0.000** | **16.793** | **0.592** |
| *Organic:salt ratio (g/g)* | *0.000* | *1.846* | *0.779* |

Simplified model matrices consisted of a roughly physiological NaCl concentration (8 g/l; ThermoFisher Scientific, 207790010) and proteins of different sizes in milli-Q water. Protein concentration ranged from 10^−9^ to 16.8 g/l, to achieve protein:NaCl dry mass ratios from ∼10^−10^:1 (corresponding to approximately one molecule of human albumin per virus) to 2.1:1 (corresponding to the total organic:NaCl ratio in SLF). Human albumin (66 kDa) and transferrin (77 kDa) were the same proteins as used in SLF, and ferritin from equine spleen (440 kDa), albumin from chicken egg white (44 kDa) and γ-globulins from human blood (160 kDa) were purchased from Sigma-Aldrich.

### Protocol for inactivation of IAV in 1-μl droplets

Prior to each experiment, purified virus stock in PBS was spiked into the matrix of interest in a 1.5-ml plastic tube (Sarstedt, 3080521), to achieve a starting concentration in the experimental solution of either 10^6^ PFU/ml (“normal titer”) or 10^7^ PFU/ml (“high titer”). The high titer experiments were introduced to expand the measurable range of inactivation. The tube was vortexed and kept in a cold tube rack until use.

Inactivation experiments in droplets were performed as described in Schaub et al.^31^ Briefly, experiments were performed in an environmental chamber (Electro-Tech Systems, 5532) with controlled RH and temperature. Experiments were conducted at 25 ± 2°C, and at the indicated RH ± 2%. Tested RH values included 15, 25, 40, 50, 60, 70, 80 and 95%.

For each virus-spiked matrix, four 1-μl droplets were deposited in four individual wells of a 96-well plate with a hydrophobic surface (Greiner Bio-One, 655901), and the last droplet was immediately collected to serve as the sample at time t = 0. The other three droplets were collected after 1 h of exposure to the RH of interest. Over the course of the experiments, the droplets were periodically monitored for efflorescence by means of visual inspection. Droplets were collected by adding 300 μl of PBSi in the well, followed by a resuspension by up and down pipetting and scratching of the bottom of the well with the pipette tip. Each sample was then separated in 2 aliquots of 150 μl and frozen at −20°C until infectivity and GC quantification. The recovering and freezing processes did not cause any further inactivation. As a control experiment in bulk, a 1-μl sample was taken directly from the IAV-spiked matrix at the beginning and at the end of the experiment and was diluted in 300 μl of PBSi. During the experiment, the plastic tube containing the spiked matrix was kept in the environmental chamber to ensure similar conditions as for the droplets. After the experiment, the chamber was disinfected using ethanol prior to opening.

Inactivation after 1 h was quantified as log(N/N_0_), where N is the number of infectious viruses in a droplet after one hour, and N_0_ is the initial number of infectious viruses in the droplet. The values were corrected for physical virus losses due to attachment to the well plate (GC/GC_0_), which was determined from the fraction of genomic copies recovered from the droplet after 1 h (GC) compared to the initial number of genomic copies in a 1-μl droplet (GC_0_).

### Statistical analysis

Ordinary one-way or two-way analysis of variance (ANOVA) combined with Tukey’s multiple comparisons test were used to compare data from groups involving a single independent variable or two independent variables, respectively. Correlations were tested by computing the Pearson correlation coefficient r. All statistical analyses were performed using GraphPad Prism v.10.0.3, allowing for an α-type error of 5%. Data below the LOQ were set to the value of LOQ/√2, according to the recommendations of Hornung and Reed,^48^ and were used as such in statistical analyses.

## Results

### Matrix composition affects IAV inactivation

IAV stability was assessed over 1 h in 1-μl droplets, at RH values ranging from 15 to 95%. The droplets consisted of three different matrices, which ranged from an organic:salt mass ratio of 0:1 (PBS) to approximately 1:1 (nasal mucus) to 2:1 (SLF) (**Table 1**). Nasal mucus and SLF droplets effloresced at already in moist air with RH ≲ 70%, whereas PBS droplets effloresced only in drier air with RH ≲ 50%. Furthermore, it took up to 20 min longer for efflorescence to occur in PBS than in SLF and nasal mucus.

Inactivation was minimal in all matrices tested at high RH (80 and 95%). In PBS, IAV stability was lowest at 50-60% RH (**Figure 1A**) and increased at lower RH. This is consistent with salt-mediated inactivation, which is expected to be the fastest in droplets just above the efflorescence RH (ERH), where exposure to highly supersaturated salt molalities is most pronounced. In SLF and nasal mucus, IAV was more stable than in PBS (ANOVA p-value < 0.0001). These matrices effloresced both at higher RH and after a shorter drying time than PBS, such that virus exposure to high salt molalities was minimized. Consequently, in SLF and nasal mucus, no more than 1-log_10_ inactivation was observed on average at any RH (**Figure 1B-C**). No significant effect of initial viral titer on IAV stability was observed (ANOVA p-value = 0.26). Controls in bulk solutions showed 0.5-log_10_ inactivation in PBS, and no measurable inactivation in the other matrices (**Figure S1**).

**Figure 1:**
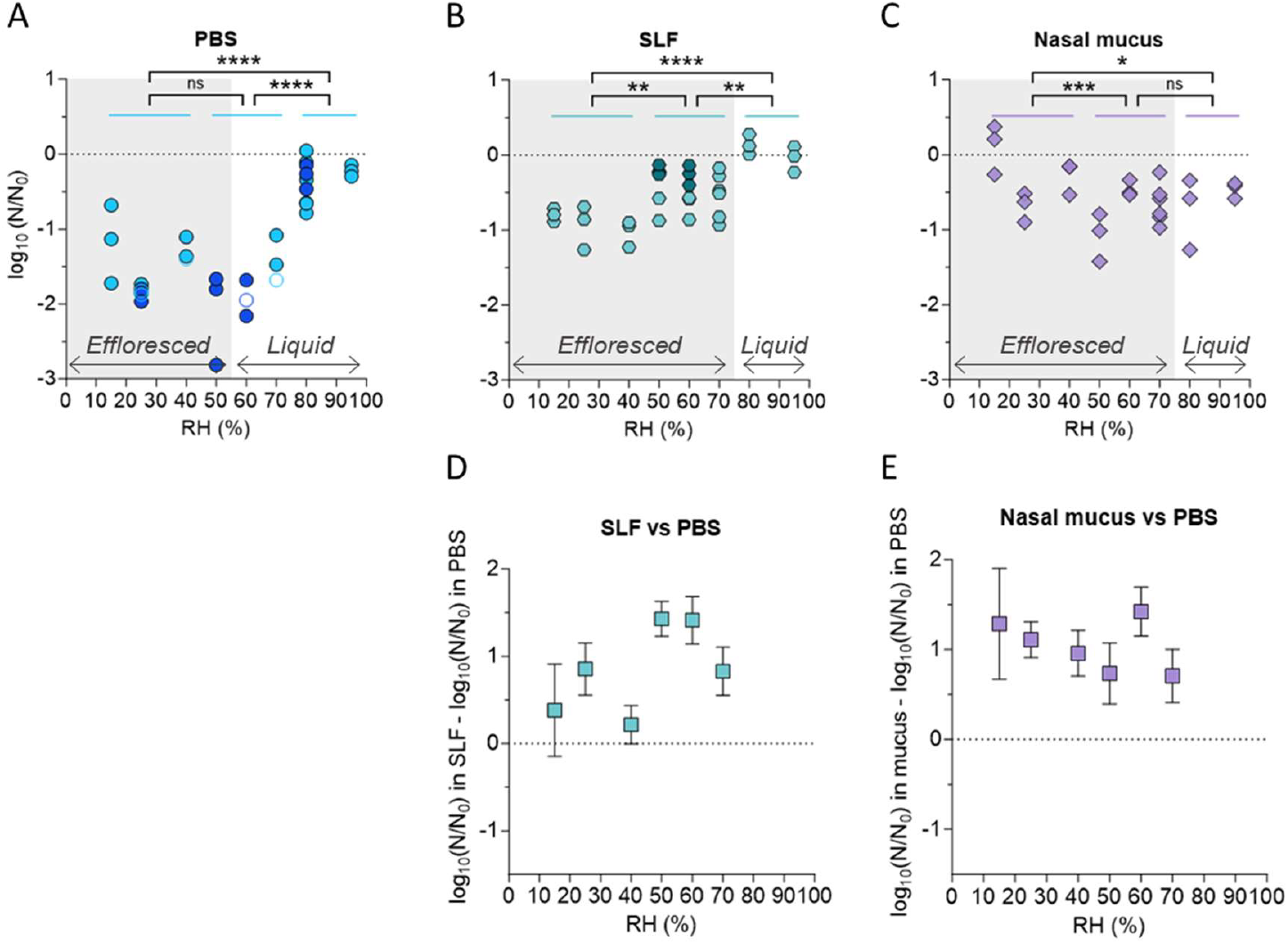
IAV inactivation after 1 h in 1-μl droplets at 15-95% RH. Droplets are composed of (A) PBS, (B) SLF, and (C) nasal mucus. Each data point represents one individual droplet. Empty symbols indicate data below the LOQ and were set to the value of LOQ/√2. The average initial titer of each droplet was 10^6^ PFU/ml (light symbols) or 10^7^ PFU/ml (darker symbols in (A) and (B)). Droplets in the gray shading were effloresced at the end of the experiment, whereas droplets outside of the shading were liquid. (D-E) Difference in inactivation between organic-laden fluids and PBS. Squares represent the average inactivation of three replicate droplets with error bars indicating standard deviations. Data were analyzed by two-way ANOVA with Tukey’s multiple comparisons test (* = p-value < 0.05, *** = p-value < 0.001, **** = p-value < 0.0001, ns = not significant).

Statistical analyses revealed a significant impact of RH on the inactivation in all matrices. Specifically, three main RH regions were compared: 15-40% RH (low RH), 50-70% RH (intermediate RH) and 80-90% RH (high RH). Inactivation levels at low and high RHs were significantly different in each individual matrix. Significant differences were also observed between low and intermediate RHs in SLF and nasal mucus, as well as between intermediate and high RHs in PBS and SLF. A detailed statistical comparison between the different RH regions is given in **Table S4**.

To determine the extent of protection provided by organics, inactivation in organic-laden matrices (SLF and nasal mucus) was compared with inactivation in PBS at the corresponding RH (**Figure 1D-E**). This analysis was only done for RH < 80%, where substantial inactivation was observed in PBS. SLF showed up to 1.5-log_10_ protection at intermediate RH (50 and 60%), while less protection was observed at lower and higher RH levels (**Figure 1D**). Mucus showed approximately 0.5-1-log_10_ protection at all RH levels < 80% RH (**Figure 1E**).

All infectivity results were corrected to account for physical virus losses due to virus adherence to the well plate. Measurements of GC/GC_0_ showed that the fraction of viruses recovered from dried droplets composed of PBS was as low as 0.01 at intermediate RHs, while the recovered fraction in SLF and mucus droplets was typically > 0.2 for all RH (**Figure S2**). This variability in virus recovery underscores the importance of accounting for physical losses when presenting infectivity data determined in deposited droplets. Failing to correct the infectivity data for this loss could introduce bias in the susceptibility of IAV to RH, especially when the recovery is low.

To assess which matrix properties impact IAV infectivity, we searched for correlations between the inactivation after 1 h (data from **Figure 1**) and the total salt and organic content of the different matrices (**Table 1**). For this analysis, we selected two RH levels, 50 and 60%, where extensive inactivation was observed in all matrices. We found poor correlations of IAV inactivation with both salt and organic content (**Figure 2A-B**). Instead, a positive trend was observed between IAV stability and the organic:salt mass ratio in each matrix (**Figure 2C**). Therefore, the organic:salt ratio seems to be a promising parameter to investigate further in the context of IAV stabilization, as opposed to absolute salt or organic content.

**Figure 2:**
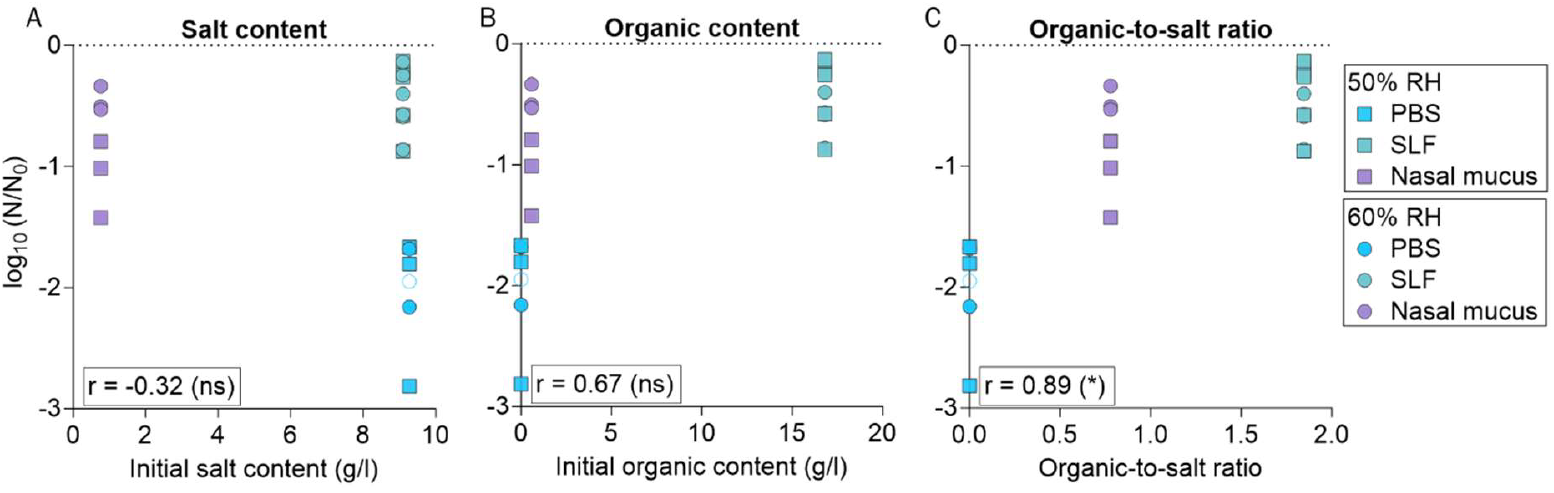
IAV inactivation after 1 h in 1-μl droplets at 50 and 60% RH as a function of salt and organic contents of PBS, SLF and nasal mucus. (A) Inactivation versus total initial salt content. (B) Inactivation versus total initial organic content. (C) Inactivation versus organic:salt dry mass ratio. The squares and circles represent inactivation at 50 and 60% RH, respectively. Each data point represents one individual droplet. Empty symbols indicate data below the LOQ and were set to the value of LOQ/√2. Also indicated is the Pearson correlation coefficient r, calculated for each panel (* = p-value < 0.05, ns = not significant). Note that for nasal mucus, other organics not quantified herein may have been present. The values of the organic content and organic-to-salt ratio of nasal mucus therefore represent minimum values.

### Proteins in SLF provide the largest IAV protection

Next, we aimed to determine which organic constituent in respiratory matrices most enhances IAV stability. To this end, we selectively removed individual organic components (either the proteins, the lipids, the antioxidants, or all three) from SLF and compared the inactivation in 1-μl droplets of these SLF derivatives to the inactivation in full SLF droplets. These experiments were performed for 1 h at 60% RH, where SLF offered the greatest protection compared to PBS (1.5-log_10_; see **Figure 1D**). The SLF derivatives were designated as “no protein”, “no lipid”, “no antioxidant”, and “only HBSS (**Figure 3**).

**Figure 3:**
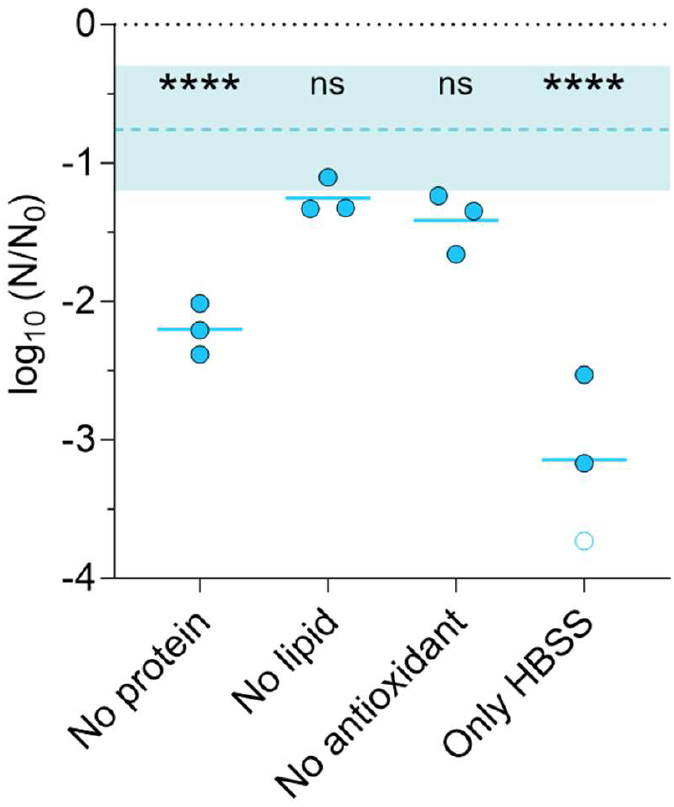
IAV inactivation after 1 h in 1-μl droplets at 60% RH in SLF derivatives. Each data point represents one individual droplet. Empty symbols indicate data below the LOQ and were set to the value of LOQ/√2. All droplets were effloresced. The shaded area indicates the range of inactivation observed in SLF droplets at 60% RH (15 replicates from 5 independent experiments) and the dashed line shows the mean. Data points below the shaded area indicate that the tested matrix is less protective than SLF; data points that fall within the shaded area indicate that protection is maintained despite the removal of the indicated component. Data were analyzed by one-way ANOVA with Tukey’s multiple comparisons test to compare the inactivation in each decomposed matrix with inactivation in SLF (**** = p-value < 0.0001, ns = not significant).

In all SLF derivatives, all droplets effloresced by the time they were sampled (1 h), though HBSS droplets effloresced later than the other SLF derivatives. The largest inactivation over 1 h was observed in HBSS only (**Figure 3**). In this solution, IAV stability was lower than in PBS (**Figure 1A**), indicating that HBSS alone offered no protective effect on IAV. When proteins were removed from SLF, the second-largest decay was observed. IAV was thus less stable than in full SLF, but more stable than in HBSS alone. Proteins are present in SLF in larger quantities than lipids and antioxidants (**Table 1**); thus, their removal leads to the greatest organic depletion. Finally, the removal of lipids or antioxidants from SLF did not significantly reduce IAV stability compared to full SLF, indicating that their contribution to IAV stabilization is minor. A detailed statistical comparison between the different samples is given in **Table S5**.

### The protein-to-salt ratio determines IAV stabilization

Human albumin is the most abundant protein in SLF, and NaCl is the main salt (**Table 1**). These constituents were therefore selected to determine how different organic (protein):salt ratios affect IAV stability in droplets. Experiments were conducted for 1 h at 60% RH (**Figure 4**). The initial NaCl concentration was 8 g/l in all experiments containing NaCl. The smallest albumin:NaCl dry mass ratio, 10^−10^:1, corresponds to approximately one albumin molecule per infectious virus. The highest tested ratio was 2.1:1, referring to an albumin concentration of 16.8 g/l and corresponding to the total organic mass in SLF (**Table 1**).

**Figure 4:**
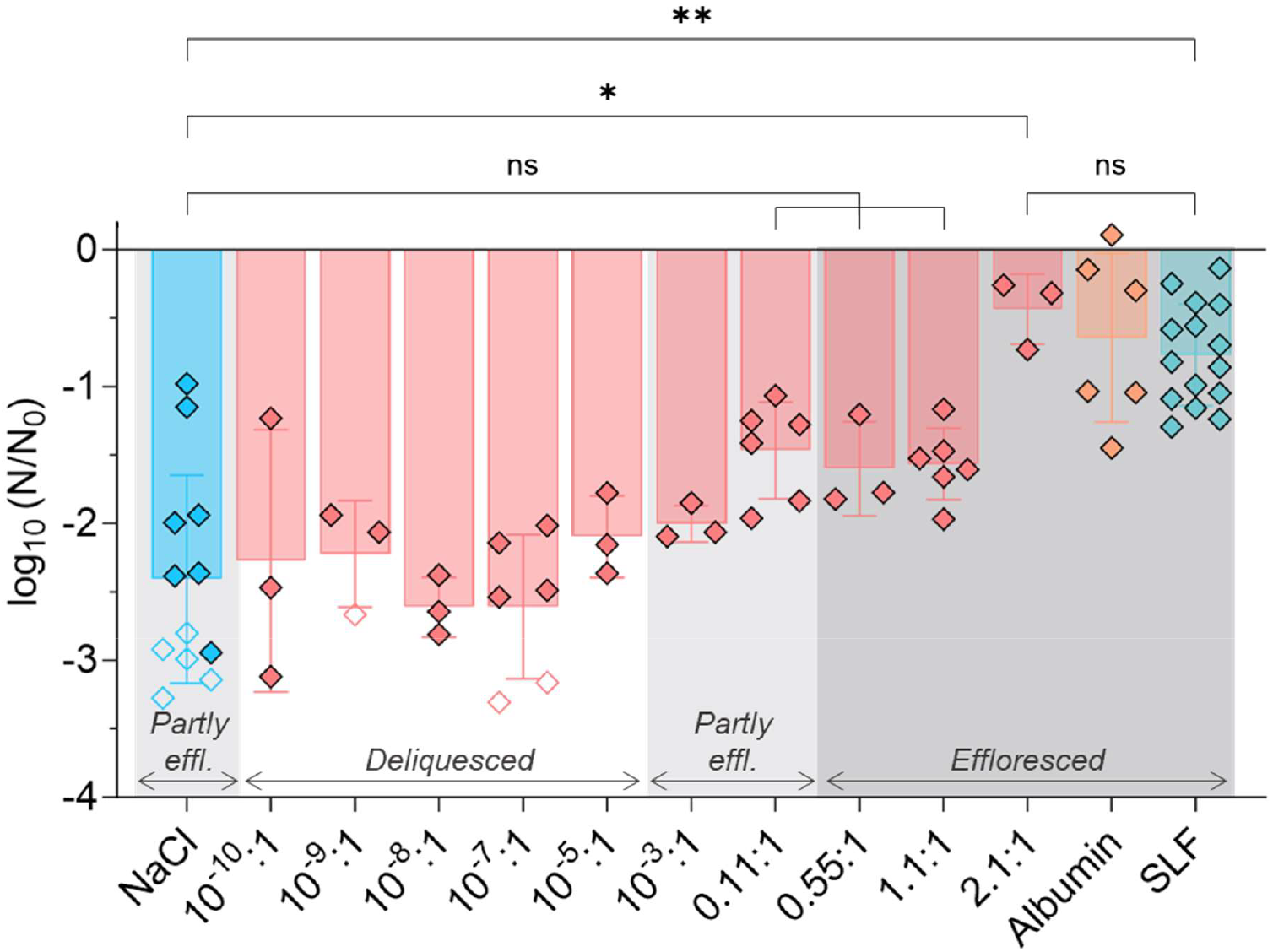
IAV inactivation after 1 h at 60% RH in 1-μl droplets composed of various albumin:NaCl dry mass ratios. The initial NaCl concentration was 8 g/l and the initial albumin concentration was varied from 10^−9^ to 16.8 g/l, according to the indicated albumin:NaCl dry mass ratio. The “albumin” droplets (without NaCl) had an initial albumin concentration of 8.8 g/l, corresponding to the albumin concentration in SLF. Each data point represents one individual droplet. Empty symbols indicate data below the LOQ and were set to the value of LOQ/√2. The mean (length of the bar) and the standard deviation (error bars) were determined based on all replicates conducted for a given condition. The dark gray shading indicates conditions where all droplets were effloresced; the light gray shading represents conditions where a subset of droplets were effloresced; and the absence of shading represents conditions where all droplets were deliquesced. Data were analyzed by one-way ANOVA with Tukey’s multiple comparisons test (* = p-value < 0.05, ** = p-value < 0.01, ns = not significant).

Even though 60% RH is above the reported ERH of aqueous NaCl (41-51%),^49^ a subset of NaCl droplets nevertheless effloresced during the experiment, possibly due to a scratch or an impurity in the well plate (e.g., dust) promoting efflorescence. Increasing amounts of albumin caused the droplets to increasingly effloresce. Droplets with albumin:NaCl ratios of 10^−10^:1 to 10^−5^:1 remained liquid over the course of the experiment. At ratios of 10^−3^:1 and 0.11:1, some droplets were effloresced and some droplets were liquid. At ratios ≥ 0.55:1, all the droplets effloresced. Albumin droplets (without NaCl; initial concentration = 8.8 g/l) presented a “coffee-ring” morphology, whereas salt-containing droplets exhibited a crystalline structure (**Figure S3**).

Consistent with the trend in efflorescence, IAV stability increased with increasing content of albumin at a constant salt concentration. Specifically, low albumin:NaCl ratios exhibited no difference in IAV stability compared to pure NaCl droplets. A protective effect of albumin of ∼1-log_10_ was first observed at an albumin:NaCl ratio of 0.11:1, corresponding to an albumin concentration of 0.88 g/l (10x lower than its concentration in SLF), though this protection was not statistically significant (p-value = 0.354). Only the matrix with an albumin:NaCl ratio of 2.1:1 and SLF were significantly more protective than NaCl only (p-value = 0.023 and 0.002, respectively). Finally, the 2.1:1 albumin:NaCl matrix, SLF and albumin only, were all similarly protective. A detailed statistical comparison between the different samples is given in **Table S6**.

### Smaller proteins are most protective

Different proteins were tested to determine whether the ability to stabilize IAV extends beyond human albumin. Transferrin, ferritin, chicken albumin, and γ-globulins were mixed with NaCl at a single protein:NaCl mass ratio of 1.1:1 (i.e., 8.8 g/l of protein and 8 g/l of NaCl). The resulting inactivation was compared to that observed in human albumin experiments at the same ratio shown in **Figure 4**. An 8-g/l NaCl solution and SLF were used separately as controls. Droplets containing IAV were deposited for 1 h at 60% RH.

Most NaCl droplets did not effloresce, while most of the other droplets effloresced after 20-35 min. An exception appeared with one of the chicken:albumin and one of the transferrin droplets that did not effloresce, indicating that 60% RH is close to the ERH of the solution.

Inactivation between 0.4 and 2.2-log_10_ was observed depending on the protein, whereas infectious virus titers in all but one NaCl droplet dropped below the LOQ. This confirms a protective effect of all proteins tested, though the extent of protection depends on the protein of interest (ANOVA p-value < 0.0001). We next considered if the protective effect can be related to the proteins’ sizes. At the same protein mass of 8.8 g/l, their molar concentrations in the droplets ranged from 0.020 mM (ferritin) to 0.20 mM (chicken albumin). When including the experimental results for human albumin:NaCl at a 2.1:1 mass ratio (**Figure 4**), the molarity range expands to 0.25 mM. We observed a statistically significant correlation (Pearson correlation coefficient r = 0.84, p-value = 0.0084) between virus stability and protein molarity (**Figure 5**), indicating that at a given mass, smaller proteins are more efficient at protecting IAV. Furthermore, this result illustrates that the molar protein-to-salt ratio, rather than the mass ratio, best describes a protein’s influence on IAV stability.

**Figure 5:**
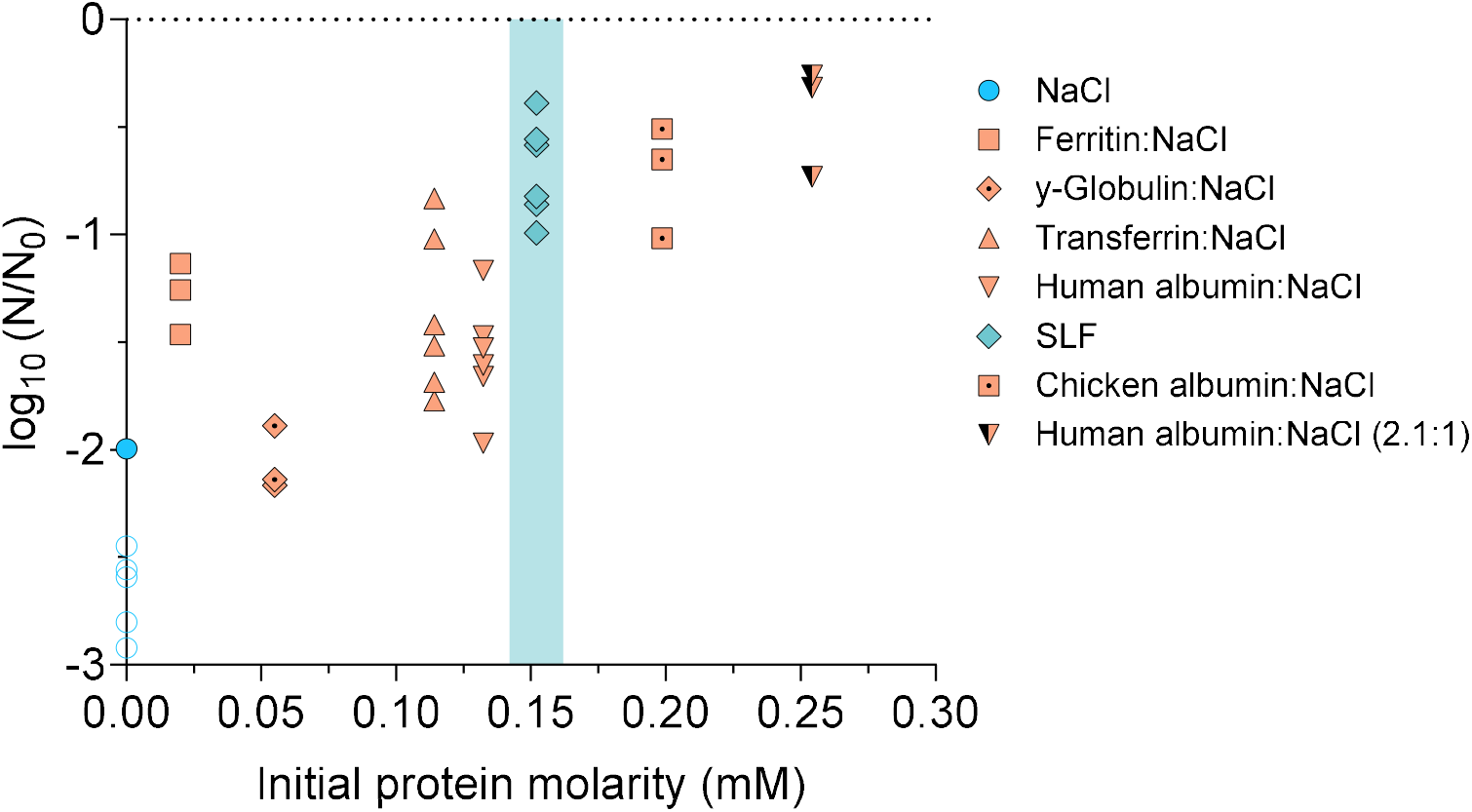
IAV inactivation after 1 h in 1-μl droplets at 60% RH in NaCl and different organics as a function of initial protein molarity. 8.8 g/l of different proteins were mixed with NaCl (8 g/l) at a 1.1:1 mass ratio. Each data point represents one individual droplet. Empty symbols indicate data below the LOQ and were set to the value of LOQ/√2. (Note that the LOQ of each data point presented depends on the fraction of viruses recovered from each individual droplet. Therefore, the LOQ varies between droplets.) All droplets were effloresced, except a subset of the NaCl droplets. The shaded area highlights SLF, which contains lipids, antioxidants and sugars in addition to proteins. Data from **Figure 4** for human albumin:NaCl at a 2.1:1 ratio are also shown.

## Discussion

IAV stability in deposited droplets has long been known to depend on RH.^18,19,30^ Our results confirm this finding, but further demonstrate that the role of RH depends on the matrix composition (**Figure 1**). We observed an increased stability of IAV when organics were present in the matrix, in particular at mid-range RH. This is in agreement with previous studies that have indicated increased stability of IAV at intermediate RH levels in the presence of organics.^18,19^ In the matrices used herein, the extent of IAV stabilization offered by organics was not associated with the absolute initial salt or organic content (**Figure 2A-B**). A similar lack of association was reported by Rockey et al.^30^ in saliva and respiratory mucus droplets. On the contrary, we observed here that IAV inactivation is inversely correlated with the organic:salt ratio of the matrix (**Figure 2C**). This indicates that this ratio is a potential determinant for virus stability in solution containing organics. Indeed, this is not surprising because inactivation increases with salt concentration and because additions of organic matter are protective for IAV, both by lowering the salt concentration and by a direct protective effect, the background of which is still poorly understood.^31^

Nasal mucus and SLF both exhibited a stabilizing effect on IAV compared to PBS, especially at intermediate RH. This may be explained by the effect of nasal mucus and SLF on salt efflorescence in droplets. In our previous work,^31^ we showed that the fastest inactivation of IAV in droplets occurs immediately before efflorescence, when NaCl molality is at its highest (supersaturated solution). After efflorescence, when NaCl molality drops to saturation, inactivation proceeds at a slower rate. The occurrence of efflorescence thus protects IAV from inactivation by reducing the virus’ exposure to supersaturated NaCl solution. The organic-laden matrices tested herein effloresced at a higher RH than PBS, such that at an RH of 60 and 70%, SLF and nasal mucus droplets are effloresced, whereas PBS droplets are deliquesced (**Figure 1**). An upward shift in ERH for SLF and nasal mucus was already observed in our previous work for smaller particles (dry radius ca. 10 μm),^16^ though the upward trend in the larger droplets used herein was more pronounced, possibly due to impurities in the matrix or imperfections on the surface. Similarly, Yang et al.^19^ reported a higher ERH in organic-rich DMEM compared to PBS. This difference in ERH may in part explain why the protective effect of SLF and – to a lesser extent – nasal mucus is particularly high around 60% RH (**Figure 1D-E**). However, protection was also high at 50% RH, when all droplets were effloresced. This can be rationalized by the finding that efflorescence in SLF and nasal mucus occurred after a shorter drying time compared to that in PBS, thus again protecting IAV from prolonged exposure to supersaturated NaCl. A similar effect was found in our work on protective effects of commensal bacteria on IAV in droplets at 40% RH: while bacteria did not affect ERH, they caused the droplet to flatten, such that water evaporation occurred more rapidly and the drying time to efflorescence decreased, thereby stabilizing IAV.^36^

The decomposition of SLF revealed that in this matrix, all organics provided some extent of IAV stabilization. The “HBSS only” droplets were the last to effloresce, suggesting that all organic components in SLF (except the dextrose already present in HBSS) can promote efflorescence and thus protect the virus from exposure to high salinity. Among the different organic components of SLF, proteins were the most protective (**Figure 3**). We did not determine whether the strong protection arises from the higher protein mass concentration compared to other organics in SLF (**Table 1**), or if proteins themselves are more efficient in stabilizing the virus. However, lipids, which were also present in substantial quantities (5.4 g/l), were not required to stabilize IAV. Furthermore, our previous work demonstrated that a similar mass concentration of sucrose (8 g/) only had a minimal protective effect on IAV stability over the course of an hour.^31^ Combined, this suggests that at similar mass, the presence of proteins is more protective than the presence of other organics.

We furthermore established that increasing the protein:salt ratio increases IAV stability, and that a threshold ratio (in this case, 0.88 g/l of albumin for 8 g/l of NaCl, corresponding to a 0.11:1 ratio) is necessary to provide stabilization of IAV by at least 1 log_10_ (**Figure 4**). In the tested matrices (SLF and nasal mucus), the protein:salt content exceeds the threshold determined for albumin, which correlates with the protection observed in SLF and nasal mucus. In comparison, Vejerano and Marr^8^ have estimated that a simulated fluid containing ∼10 g/l of salts, ∼10 g/l of proteins and ∼0.6 g/l of surfactant could accurately model the composition of respiratory fluid, while Niazi et al.^50^ proposed a fluid containing 10 g/l of salts and 8 g/l of proteins. The protein:salt ratio in such fluids also meets the threshold necessary for virus protection, suggesting that these fluids would also induce virus stabilization in drying respiratory droplets.

The mechanism underlying the increasing IAV stabilization with increasing albumin content may again be linked to the effect of the protein on salt efflorescence. Droplets with a low human albumin content (albumin:NaCl ≤ 10^−5^:1) always remained deliquesced and exhibited > 2 log_10_ inactivation, whereas droplets with albumin:NaCl ratios ≥ 0.11:1 were partly or fully effloresced and exhibited lower levels of inactivation. In addition, albumin droplets exhibited a “coffee-ring” (**Figure S3**), which coincided with high stability for the virus, consistent with the protection mechanism proposed by Huang et al.,^34^ which suggests that the aggregation of viruses and proteins within this coffee ring prevents inactivation.

Proteins other than human albumin were also found to stabilize IAV (**Figure 5**), but their effect differed both on a mass and on a concentration basis. Specifically, the smallest protein, chicken albumin, provided the largest protective effect. The observed IAV stabilization can only partly be explained by the proteins’ role in promoting efflorescence. Efflorescence in all protein-containing droplets occurred earlier than in NaCl alone, which explains the stabilization of IAV compared to pure NaCl droplets. However, in droplets containing chicken albumin, efflorescence occurred concurrently with droplets containing other proteins, yet a greater stabilization of IAV was observed. Therefore, the various levels of protection provided by the different proteins cannot be explained by differences in promoting efflorescence. Instead, protection by proteins was found to be correlated with protein molarity, which – at a given protein mass – is inversely correlated to protein size. Smaller proteins are thus more efficient at stabilizing IAV. This effect may arise from potential differences in the morphology of the dried droplets, or with differing affinities of IAV to the proteins tested. Rockey et al.^30^ observed that the matrix composition modulates the morphology of the dried droplets, and this morphology modulates in turn the infectivity of IAV. Similarly, Huang et al.^34^ observed increasing coffee ring effect with increasing initial protein concentration, suggesting that a higher initial protein molar content may stabilize the virus more efficiently. Whether the high protein molarity in the droplets containing small proteins indeed led to a more significant coffee ring remains to be assessed in future microscopy studies.

This study has a number of limitations. First, we assessed virus inactivation after only a single time period (1 h) of exposure at a given RH. This time reflects a realistic exposure time to exhaled viruses in a class room, gym or on public transportation. However, more detailed kinetic experiments would be beneficial to determine the inactivation dynamics and to identify the stage of the drying process at which organics are most protective. In addition, a full mechanistic explanation for the protection provided by organics is still lacking, though our data suggest that the promotion of efflorescence by organics can partly account for the observed protective effects. Finally, while we were able to confirm the protective effect of organics in deposited droplets on a specific surface, additional work in an airborne particles system is required to establish the IAV stabilizing effect in the smallest particles exhaled by infected individuals.

In summary, we demonstrate that the addition of organic components in drying droplets enhances the stability of IAV and limits its inactivation. The most protective components in SLF are the proteins, which are also the most abundant organics. We identified two factors that promote virus stability in respiratory matrices: first, IAV is stabilized in matrices with a protein:salt ratio that is sufficiently high to promote efflorescence by increasing the ERH and accelerating the onset of efflorescence at a given RH (here protein:salt ≥ 0.11); and second, small proteins that result in a high molar protein concentration are more efficient at stabilizing IAV. The understanding of the role of organics in the stabilization of airborne viruses is essential to comprehend the mechanism underlying virus transmission through exhaled aerosol particles and droplets.

## ASSOCIATED CONTENT

Supporting information

The following file is available free of charge and contains:

Method descriptions and composition tables for analysis of lipids, main ions and proteins in nasal mucus; controls and recovery fractions for droplet experiments; droplet pictures; check-list for RT-qPCR following MIQE guidelines; details for ANOVA analyses (PDF)

## AUTHOR INFORMATION

### Author Contributions

Conceptualization: AS, SCD, TK

Data curation: AS

Formal analysis: AS

Methodology: AS, TK

Investigation: AS, IG, KV

Resources: SSt, TK

Writing–original draft: AS, TK

Writing–review & editing: AS, BL, CT, GM, IG, KV, LKK, MP, NB, SCD, SSt, TP, TK, UKK

Visualization: AS

Supervision: SSt, TK

Project administration: TK

Funding acquisition: AN, SSt, TK, TP, UKK, WH

### Notes

Authors declare that they have no competing interests.

### Data availability

Experimental data are available in the public repository Zenodo under the following link: https://zenodo.org/records/11204219.

## ACKNOWLEDGEMENTS

This work was funded by the Swiss National Science Foundation (grant 189939).

